# APICE-Py: An Open-Source MNE-Python Pipeline for Scalable EEG Preprocessing

**DOI:** 10.64898/2026.07.23.740250

**Authors:** Nicolò Formento Moletta, Ana Fló, Ghislaine Dehaene-Lambertz, Jhunlyn Lorenzo

## Abstract

Electroencephalography (EEG) is fundamental to cognitive neuroscience as it provides a direct measure of human neural activities with millisecond precision. Its noninvasive nature allows for the study of brain function across diverse age groups and experimental contexts—from newborns to adults, and from tightly controlled laboratory environments to more naturalistic real-world settings. However, EEG signals—especially those recorded from infants—are highly prone to noise and artifacts, posing significant challenges for data analysis. To address these issues, we present APICE-Py (Automated Preprocessing for Infants Continuous EEG), an open-source preprocessing pipeline originally designed as a matlab toolbox for infant EEG and now re-implemented in Python to support scalable and flexible analysis across developmental and adult datasets. APICE-Py is built upon three core principles: (i) adaptive artifact detection on continuous data using data-driven thresholds rather than fixed cutoffs; (ii) hierarchical artifact correction, combining short-segment correction via Principal Component Analysis (PCA) with broader segment and continuous data correction using Spherical Spline Interpolation (SSI); and (iii)transparent reporting, providing comprehensive quality logs and decision-tracking to ensure reproducibility and informed analysis. We summarize the underlying algorithms, release an implementation compatible with common EEG formats, and demonstrate its use on three datasets spanning neonates, 5-month-old infants, and child–parent hyperscanned data, acquired using high-density wet electrodes and mobile gel-based EEG systems. When benchmarked against the original MATLAB implementation, APICE-Py achieved comparable levels of data quality and trial retention. While the original pipeline was developed for early developmental EEG (e.g., infants), we show that the pipeline further extends its applicability to both children and adult datasets, enabling robust preprocessing across a broad age range. Moreover, it supports data acquired using a variety of EEG configurations and experimental settings, highlighting its flexibility across age groups, hardware systems, and paradigms. The APICE-Py source code and documentation are freely available at https://github.com/neurokidslab/apice-py.

## Introduction

By capturing rich neural information with high temporal resolution, EEG has become an essential imaging modality for studying the human brain. It offers a safe and noninvasive method for measuring neural activity across a wide range of age groups, from newborns and infants to children and adults. Compared with other neuroimaging techniques, such as magnetic resonance imaging (MRI) and magnetoencephalography (MEG), EEG is more tolerant to movement, allowing its use across a wide range of experimental paradigms—from resting-state conditions and cognitive tasks to tasks involving naturalistic behaviors such as locomotion, object manipulation, and social interaction—as well as across diverse environments, including hospitals, laboratories, and real-world settings. In addition, EEG allows the collection of large-scale data over extended periods and, with hyperscanning technology, enables simultaneous neural recordings from multiple individuals, thereby facilitating the study of dynamic brain activity and interindividual variability.

Despite its many advantages, EEG data are highly susceptible to various sources of noise and artifacts that can generate electrical potentials up to ten times larger than the underlying brain signals (Delorme (2023), Hervé et al. (2022)). Non-neural activities such as ocular movements, facial muscle contractions, heartbeats, respirations, and body movements inevitably distort the EEG signals. While adults can be instructed to minimize body movements and focus on the task, this level of behavioral control is challenging with young participants. Moreover, neural signals and their power spectra vary considerably across developmental stages due to brain maturation, making it particularly difficult to distinguish true neural signal from slow drifts in infants EEG compared to adults. The choice of EEG system also plays a critical role in data acquisition and analysis (Chaddad et al. (2023)). High-density systems, which facilitate more accurate signal reconstruction due to their larger number of electrodes, are typically equipped with wet sensors that can exhibit lower signal-to-noise ratio (SNR). In contrast, gel-based electrodes typically provide higher SNR and greater signal stability over time, owing to better scalp contact and lower impedance. However, these systems often use fewer electrodes (64 and often less in pediatric populations), limiting spatial resolution and reducing signal reconstruction accuracy. Environmental conditions can also contaminate EEG recordings through line noise (50/60 Hz), inadequate electrical shielding, cable movement, or the physical proximity of another person to the participant. These factors can significantly degrade signal quality and complicate the extraction of pertinent neural information.

To address these challenges, our team developed an automated preprocessing pipeline called APICE (Fló et al. (2022)), designed to remove artifacts and reconstruct EEG signals. Good statistical power requires as many good-quality trials as possible. While it is preferable to acquire high-quality data, rather than relying on extensive preprocessing (Luck (2014); Delorme (2023)), EEG acquisitions remains challenging in many situations. For instance, children and patients tend to have shorter attention spans and lower task compliance compared to healthy adults, making it difficult to obtain a sufficiently large number of artifact-free trials. Consequently, it is essential to preserve as much usable data as possible while minimizing the degree of data reconstruction. Attrition rates are also higher in younger participants (Hervé et al. (2022)), as recordings are often interrupted by movement, fussiness, or loss of attention. Although this data loss could theoretically be mitigated by recruiting larger samples, this approach poses practical and logistical constraints, even in adult populations. Thus, EEG research often involves a trade-off between sample size and data quality.

Traditionally, multiple researchers would visually inspect EEG data to manually identify artifacts and noisy segments or channels, with inter-rater agreement used to define the final cleaned dataset (Shirk et al. (2017)). Fixed thresholds have also been applied to reject abnormally high amplitudes, while auxiliary recordings such as the electrooculogram (EOG) and electrocardiogram (ECG) helped identify blink and heartbeat components, and synchronized videos were used to annotate participant movements (Hervé et al. (2022)). However, these procedures are time-consuming, subjective, and impractical for large datasets, and they often fail to account for variability across participants, age groups, and experimental designs. To overcome these limitations, several standardized preprocessing pipelines have been developed to more effectively manage signal noise, artifact correction, and the requirements of subsequent analyses (Mumtaz et al. (2021), Hervé et al. (2022), Delorme (2023), Coelli et al. (2024), Kumaravel et al. (2022)). Early standardized pipelines such as PREP, FASTER, MARA, and Automagic were primarily optimized for adult EEG, whereas newer frameworks, including HAPPE, MADE, EEG-IP-L, EPOS, APICE, iMARA, and NEAR, were specifically designed for developmental data. Most pipelines, aside from the standard APICE, PREP, and NEAR, rely on Independent Component Analysis (ICA) to identify and remove artifacts such as eye blinks and cardiac activity; however, determining which and how many components to reject remains subjective. In low-density EEG systems, where multiple neural sources may be mixed within a single component, component rejection can inadvertently eliminate genuine neural signals. Although ICA performs robustly in adult EEG, its application to infant data is challenging due to greater signal variability and the prevalence of slow-wave activity (Hervé et al. (2022)). Several pipelines (excluding APICE, iMARA, NEAR and EEG-IP-L) also rely on fixed amplitude or variance thresholds for artifact detection, which fail to account for individual and age-related differences as well as changes in EEG background activity associated with different stages of vigilance. Consequently, these methods may either over-reject clean data or retain residual noise. Other frameworks incorporate machine-learning tools that require model training (e.g., MARA, iMARA) or statistical techniques such as Artifact Subspace Reconstruction (ASR) that depend on short calibration periods (NEAR, PREP). While such methods are feasible for adult EEG, they are less practical for infant recordings, which are typically shorter, noisier, and contain fewer clean segments for reliable calibration or training (Chaddad et al. (2023)). In contrast, pipelines implementing subject-specific or adaptive thresholds, such as APICE and EEG-IP-L, can better account for inter-individual and developmental variability. Adaptive approaches dynamically adjust detection limits based on the statistical properties of each recording, enabling more accurate artifact identification while preserving a larger proportion of usable data (Fló et al. (2022)). Using adaptive thresholds is particularly advantageous when processing large datasets, as the pipeline parameters are automatically adjusted and artifact rejection is fully automated. This contrasts with ICA-based approaches, which require manual inspection and decision-making for each component, making them less scalable and more operator-dependent (Hervé et al. (2022)).

In this paper, we introduce APICE-Py, a Python-based implementation of the Automated Preprocessing of Infants Continuous EEG (APICE) pipeline. Our objectives were twofold: (1) to provide an open-source, scalable, and reproducible framework for EEG preprocessing that extends the functionality of the original MATLAB version into the MNE-Python ecosystem; and (2) to demonstrate that, beyond its developmental focus, APICE-Py can robustly process EEG data across diverse age groups, electrode configurations, and experimental paradigms, thereby enhancing its applicability to both research and clinical settings. APICE-Py features (i) automatic artifact detection using adaptive, data-driven thresholds; (ii) hierarchical correction methods for reconstructing short-duration artifacts and maximizing data retention; and (iii) transparent quality metrics and reports that promote reproducibility and data sharing. Alongside the source code, we provide a tutorial to guide users in customizing APICE-Py for their specific research needs, in line with the principles of open science (Buzzell et al. (2023), Coelli et al. (2024)).

### Pipeline Overview

We developed APICE-Py as a Python implementation of the original MATLAB/EEGLAB-based APICE pipeline (Fló et al. (2022)) and built upon the MNE-Python framework (Gramfort et al. (2013)). Figure 1 illustrates the default APICE-Py preprocessing workflow. The pipeline is fully modular, allowing users to adapt or extend individual steps to meet specific research and analysis needs using the functions provided in the APICE-Py module. Depending on the analysis requirements, APICE-Py can process continuous EEG data or epochs for event-related potential (ERP) and other time-locked analyses. We recommend performing artifact detection and correction in two successive stages, first on continuous data, then on epoched data, to increase flexibility. Processing at the continuous stage provides an initial screening that can be saved, allowing the second stage to be flexibly adapted for different analyses of the same recordings, whether by varying epoch time windows or onset markers (e.g., when sequences of stimuli are chained), or by adjusting noise criteria depending on the analysis type, without the need to repeat the procedure on the entire dataset.

**Figure 1.**
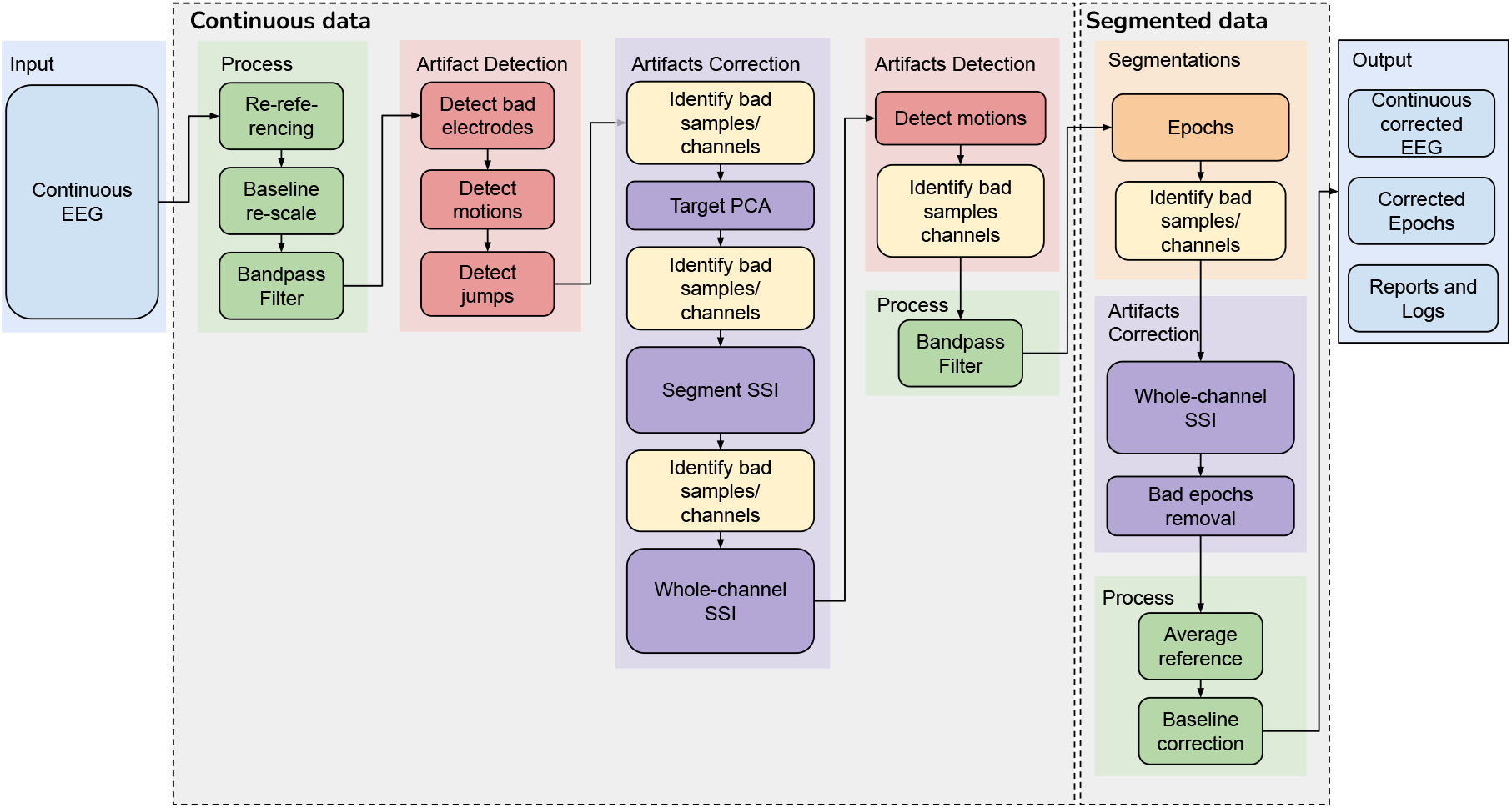
Overview of the APICE-Py preprocessing pipeline.

The first stage of the APICE-Py pipeline focuses on continuous data preprocessing and artifact handling. APICE-Py supports input data in .fif, .set, and .mff formats, as well as any other file types compatible with MNE. It also includes functions to convert ‘stim’ channels into events, define electrode layouts, and perform other data preparation steps. The raw continuous EEG data then undergo fundamental preprocessing operations: demeaning all electrodes to minimize common noise, baseline correction to stabilize the signal, and band-pass filtering within the desired frequency range (typically 0.2–40 Hz, or just below the power-line frequency). Filtering is performed using MNE’s overlap Finite Impulse Response (FIR) filter, which processes the signals in segments and recombines them to produce the final output (Gramfort et al. (2013)). Artifact rejection in APICE-Py is performed through a series of algorithms designed to identify and flag contaminated data segments. Bad electrodes—typically characterized by high impedance or persistent noise—are detected based on signal amplitude in different frequency bands, and low correlations with the other channels, as done in other pipelines, such as in PREP (Bigdely-Shamlo et al. (2015)). Motion artifacts are identified using amplitude- and variance-based measures, including deviations in running averages, while abrupt signal discontinuities or “jumps” are detected through algorithms sensitive to sudden changes in signal dynamics. In the default configuration, an initial fixed threshold of 500 µV is applied to flag excessively high-amplitude signals. Although this value is relatively high, it serves as a preliminary screening step, since subsequent adaptive algorithms more precisely identify and reject high-amplitude noise. These adaptive algorithms compute channel- and data-specific thresholds according to the Tukey criterion *Thresh*_+_ = *Q*_3_ + *k IQR* and/or *Thresh* = *Q*_1_ −*k IQR*, where the constant coefficient *k* = 3 by default, and the interquartile range *IQR* = *Q*_3_ −*Q*_1_. This approach ensures that artifact detection scales dynamically with the statistical properties of each recording.

Once artifacts are detected, APICE-Py records their locations in a Bad Channel and Time (BCT) logical matrix of size (*n*_*channels*_ × *n*_*timepoints*_). From this matrix, two summary metrics are derived: Bad Channels (BC) — a vector of size (1 × *n*_*channels*_) — and Bad Times (BT) — a vector of size (1 × *n*_*timepoints*_). A channel is classified as bad if more than 50% of its samples are flagged as contaminated, whereas a timepoint is considered bad if a substantial proportion of channels (e.g., >30%) are marked as artifacted. The identification of bad samples and channels is then repeated after each detection and correction step to ensure consistent data quality. These default percentages can be easily changed for specific needs.

Artifact correction in APICE-Py follows a three-stage hierarchical interpolation approach applied sequentially to individual channels, short segments, and finally, the entire continuous (or epoched) dataset. In the first stage, PCA is applied limited to the short artifacted segments (e.g., lasting less than 100 ms) and the components carrying most of the variance (e.g., 98%) are removed. Since PCA is limited to segments containing brief artifacts we called it target PCA. The reconstructed segments are then aligned to the adjacent clean samples. Next, SSI is applied to short segments based on spatial information from neighboring channels. In the final stage, SSI is applied to the entire continuous data to correct channels that remain globally bad throughout the recording. By default, a bad channel within a segment or the whole continuous data is interpolated only if at least 50% of its neighboring channels are classified as good. All corrected samples are logged in the Corrected Channels and Times (CCT) matrix, ensuring transparent tracking of modifications. This hierarchical correction strategy enables robust artifact removal while preserving overall signal integrity.

Following artifact correction, the data undergo an additional motion-artifact detection step to identify residual noise or interpolation-induced signal discontinuities, thereby ensuring the reliability of the cleaned dataset. For subsequent analyses, such as event-related potential (ERP) studies, an optional secondary band-pass filter (default: 0.2–20 Hz) can be applied to further refine the signal bandwidth. This step is particularly important for epoch-based analyses or other applications that require enhanced frequency specificity.

Stage 2 focuses on epoched-data. Once the continuous EEG signal is divided into analysis-ready epochs, an epoch-level artifact detection and correction using SSI are performed. Bad epochs are systematically identified and excluded or flagged based on three strict quality criteria: (i) temporal integrity — no bad time points within the epoch; (ii) correction limit — no more than 50% of the samples interpolated; and (iii) channel integrity — no more than 30% of channels marked as bad. The corrected and segmented data are then re-referenced using an average reference (applied only when the dataset contains 30 or more EEG channels) and subsequently baseline-corrected.

APICE-Py outputs both corrected continuous and segmented datasets, with annotations marking residual artifacts and interpolated segments. A comprehensive log file is also generated, detailing all preprocessing steps, detected artifact types (e.g., motion, jumps, bad channels), and the corresponding corrective actions. For transparency, users can review this log to trace the pipeline operations and identify dominant noise sources. In the Python version, APICE-Py additionally provides an interactive HTML report summarizing key data quality metrics, including artifact distribution and channel status. Data are saved following the MNE data structure: preprocessed continuous files retain artifact annotations that specify the onset, duration, and nature of uncorrected or excessively noisy segments, while epoched data include artifact information stored within the *epochs.comments* and *epochs.metadata* fields.

Lastly, APICE-Py supports batch preprocessing, facilitated by its adaptive artifact-detection framework, allowing large datasets to be processed automatically with minimal user intervention. The full source code is freely available on GitHub (https://github.com/neurokidslab/apice-py), along with a tutorial that guides users in customizing the pipeline for their specific research needs.

## Methods

Because there is no universally accepted ground-truth EEG signal, preprocessing outcomes often vary across researchers, laboratories, and software environments. Even widely used toolboxes such as EEGLAB, MNE, and Brainstorm can produce different results despite pursuing the same analytical objectives (Delorme (2023)). To establish a consistent reference point, we benchmarked APICE-Py against the original MATLAB implementation using multiple datasets spanning neonates to adults. The MATLAB version of APICE has been previously validated on several developmental EEG datasets (Fló et al. (2022); Hervé et al. (2022)), demonstrating reliable artifact detection and data reconstruction performance. Using it as a reference thus provides a meaningful standard for assessing the fidelity and consistency of the Python version. The evaluation compared data quality, artifact-rejection efficiency, signal preservation, and processing time across multiple quantitative metrics.

### A. Datasets Description

To assess the robustness and flexibility of APICE-Py, we tested it on three representative EEG datasets encompassing a wide developmental range—from neonates to adults—as well as two different recording systems, including a low- and a high-density montages (32–128 channels) with wet and gel-based electrode types, acquired under different experimental paradigms.

- Dataset 1 – Auditory task with neonates: Neural activity from 24 healthy full-term infants (11 males) was recorded during sleep or quiet rest in a sound-proof booth at Port Royal Maternity (AP-HP, Paris, France). Each neonate was presented with 216 trials consisting of four or five 250-ms syllables delivered every 600 ms. EEG was acquired using a 128-electrode EGI (Electrical Geodesics, Inc.) HydroCel net (vertex reference; 250 Hz sampling frequency).
- Dataset 2 – Visual association task with 5-month-old infants: This dataset (N = 26; 12 males; mean ± SD = 22.98 ± 1.41 weeks) corresponded to a visual association task. Each trial began with a 0.6-s central attention grabber, followed by a 1-s image, then a second 1.2-s image, and finally another 1.0–1.2-s attention grabber. Testing was terminated when infants showed signs of fussiness, resulting in an average of 126.35 ± 26.07 valid trials per participant. EEG was recorded using a 128-channel EGI HydroCel net (vertex reference; 500 Hz sampling rate) while infants sat on their parents’ laps inside a shielded booth at NeuroSpin (Gif-sur-Yvette, France).
- Dataset 3 – Collaborative task with (a) 7-year-old children and (b) their parents: This hyperscanning EEG dataset included 31 child–parent pairs recorded simultaneously using synchronized 32-channel gel-based Smarting Pro systems (mBrainTrain; 500 Hz sampling rate). Each dyad performed a number-puzzle task inside a shielded booth at NeuroSpin (Gif-sur-Yvette, France), under naturalistic interaction conditions that allowed free movement and verbal communication. Each pair completed three trials, with a mean duration of 5.6 ± 2.07 minutes.

All participants (or the parents or legal guardians of minors) provided written informed consent in accordance with institutional ethical approval.

### B. Evaluation Criteria

All datasets were preprocessed using both the APICE (MATLAB) and APICE-Py (Python) pipelines with identical parameter settings. The performance of the two implementations was then compared across the following quantitative metrics.

#### B.1. Standardized measurement error (SME)

For Datasets 1 and 2, we preprocessed and segmented the continuous EEG data into epochs to compute ERPs for each participant. SME serves as a robust index of EEG data quality and precision (Luck et al. (2021)). It quantifies the variability of ERP components (e.g., mean amplitude or latency) across trials or participants, where lower SME values indicate higher data quality and consistency. To compute a participant’s SME, the averaged ERP scores were first obtained within a defined time window of interest (e.g., peak amplitude or mean voltage). To increase the number of observations without additional data collection, we applied a bootstrapping procedure by randomly resampling n responses (with replacement) from the available n trials per subject, repeating this process 1000 times. The SME for each participant was then estimated as:

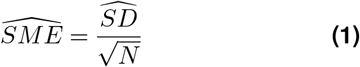

where 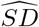 denotes the within-subject standard deviation of the bootstrapped ERP scores, and *N* is the number of samples. Finally, grand-average ERPs were computed across participants for each condition to provide a consolidated measure of the neural response to the auditory and visual stimuli.

#### B.2. Signal-to-Noise ratio (SNR)

Dataset 3 represents a continuous, interactive task. To assess data quality in the time domain, we computed the SNR of the recorded EEG signals. The raw EEG data were initially band-pass filtered between 0.2 and 40 Hz and preprocessed. The signals were then segmented into 1-minute windows with a 10-second overlap. For zero-mean signals, the SNR (in dB) was calculated as follows:

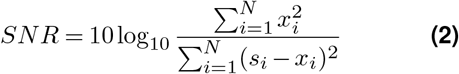

where *x*_*i*_ denotes the noise-reduced signal (artifacts were detected and corrected) and *s*_*i*_ represents the bandpass filtered signal at time *i* (Radüntz (2018)). Mean SNR values were then computed across all channels and participants, along with the mean percentage of artifact-contaminated data segments.

#### B.3. Processing time

To assess the computational time of the SSI step between the two implementations, we used a sample from Dataset 2 (128 channels x 250 seconds). Here, the raw data were considered as artifact-free, after which simulated artifacts were introduced by randomly tagging segments from a subset of electrodes as “artifact”. The proportion of affected electrodes was progressively increased from 10% to 50%, with each artifacted electrode containing between 1 and 10 artifacted segments. Each segment had a duration between 0.5 and 10 seconds. This procedure generated diverse test scenarios to provide a broad estimate of computational efficiency. The same BCT matrices were used for both implementations to ensure comparability. Computation time was measured as the total wall-clock duration required to execute the complete interpolation process under each condition.

## Results and Discussion

In the context of APICE-Py, scalability refers to the pipeline’s capacity to efficiently handle increasing complexity in both data and usage. This includes flexibility across datasets of varying sizes and recording conditions—from short to long sessions, low-to high-density montages, and infant to adult EEG—as well as computational efficiency when processing large datasets. It also encompasses the pipeline’s modular design and seamless integration with the MNE-Python framework, which enables users to adapt or extend individual components according to their analytical needs. Finally, APICE-Py is scalable at the user level, being accessible to individual researchers while supporting broader adoption within collaborative and open-science contexts.

We developed APICE-Py following the principles of APICE (Fló et al. (2022)), combining fixed and data-driven adaptive thresholds for artifact detection with hierarchical correction steps. Figure 2 illustrates an example of artifact detection and correction on a sample EEG recording (Dataset 3a; 32 channels x 250 s; 500 Hz sampling rate). The artifacts maps derived from the BCT matrices show the distribution of artifacts before (Fig. 2a) and after (Fig. 2b) correction, plotted using *plot*_*artifact*_*structure*(). In the artifact map, nonfunctional channels appear in yellow, globally noisy periods in purple, and localized contaminated segments in red. After correction (Fig. 2b), bad channels are interpolated, and most noisy segments are successfully reconstructed. The remaining bad data correspond to segments that were too long for target PCA correction and had insufficient neighboring electrodes for reliable interpolation, while the remaining bad time indicates periods with too many electrodes with remaining bad data. The superimposed original and corrected EEG traces in Fig. 2c, demonstrate APICE-Py’s efficiency to correct high-amplitude transients and sudden discontinuities. Then, Fp2 was flagged as a bad channel due to its weak amplitude correlation with neighboring electrodes. The accompanying topographic maps (Fig. 2d–e), plotted using *plot*_*percentage*_*of*_*bad*_*data*_*across*_*sensors*(), display the proportion of bad data per electrode, demonstrating a clear reduction in artifact contamination.

**Figure 2.**
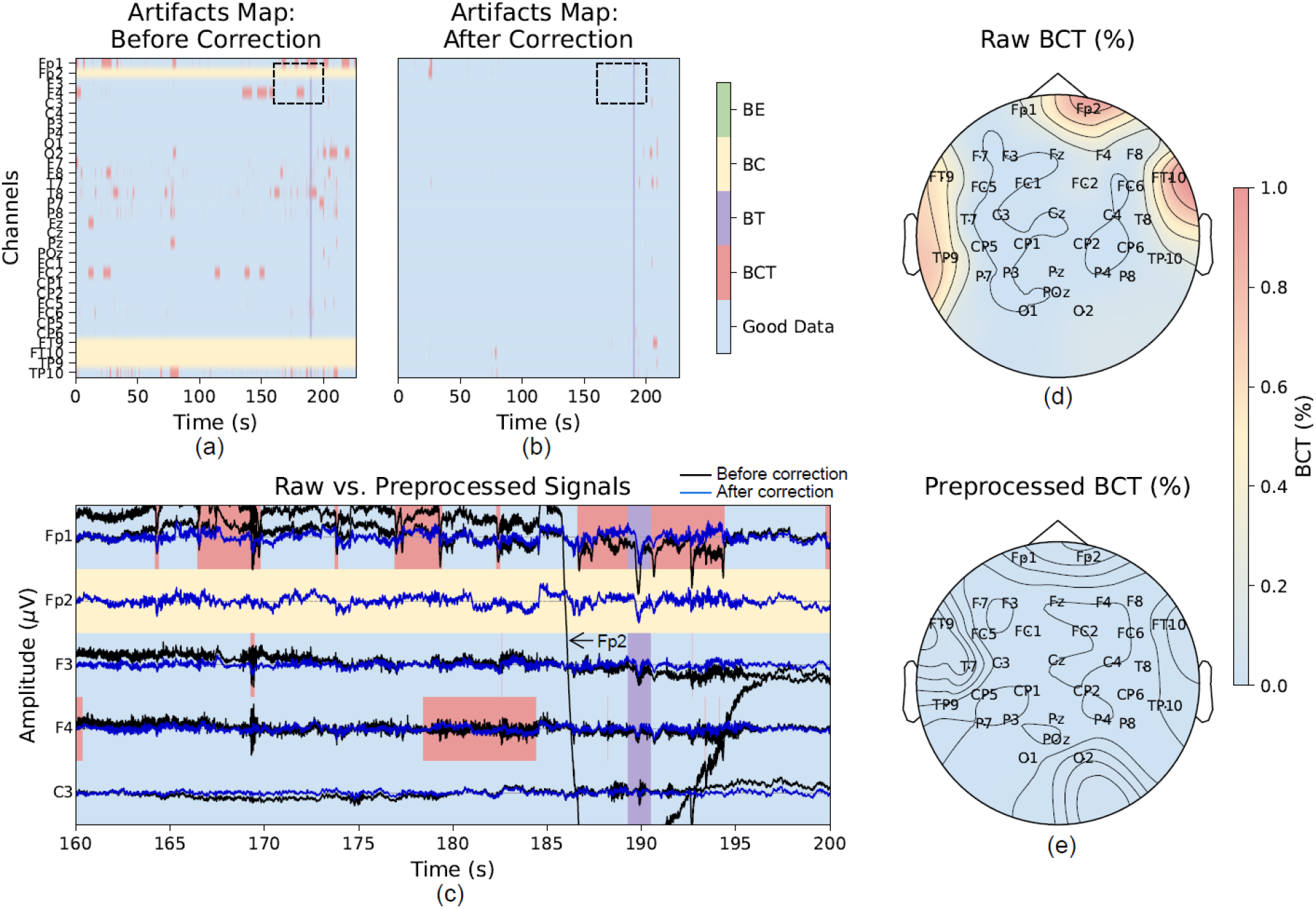
Example output from the APICE-Py pipeline applied to a sample child EEG recording. (a) Artifacts map before correction, (b) Artifacts map after correction of recoverable artifacts, (c) Excerpt of the artifact map before correction from (a) superimposed on the EEG trace. The original artifact-contaminated signal is shown in black, and the corrected signal in blue. FP2 has been identified as a bad channel, thus is highlighted in yellow, while red rectangles indicate transient bad samples within a specific channel (labeled BCT in panel a) and (d–e) topographic maps showing the proportion of artifacted data per electrode before and after correction.

These maps show that APICE-Py successfully detected and corrected artifacted channels (Fp2, FT9, FT10, and TP9). In addition, APICE-Py automatically generates an interactive HTML report that visualizes the data at each preprocessing stage, including power spectra, detected artifacts, corrected signals, ERP images, time courses, and topographic maps. Together, these outputs provide a transparent overview of the preprocessing workflow and data quality.

### C. Data Retention

To compare the artifact-detection and correction performance of APICE and APICE-Py, we preprocessed raw EEG data from four datasets using the default pipeline (k = 3) and identical parameter settings for both implementations. Differences between the two versions were assessed using paired t-tests or Wilcoxon signed-rank tests, depending on the normality of the distributions. Overall, APICE-Py detected a higher proportion of artifacts than the MATLAB implementation. This effect was most evident in the BCT metric for Dataset 2 (Python: 30.09 ± 13.62%; MATLAB: 25.81 ± 10.58%; p = 0.005**) and Dataset 3a (Python: 10.52 ± 14.46%; MATLAB: 8.28 ± 5.33%; p = 0.025*), as shown in Fig. 3a. In addition, the Python implementation detected a higher number of bad channels (BC) in Dataset 2, further indicating increased sensitivity to artifact-related deviations. Because the proportion of detected artifacts differed between implementations, the percentage of remaining artifacts after correction also showed corresponding differences between APICE and APICE-Py. Figure 3 highlights two key observations. First, EEG signal properties vary substantially across age groups, from sleeping neonates (Dataset 1) to freely moving adults (Dataset 3b). Second, despite this variability, both APICE and APICE-Py successfully preprocessed the recordings and preserved most of the usable data, demonstrating that the pipeline robustly accommodates a wide range of developmental stages, electrode configurations, and experimental paradigms.

**Figure 3.**
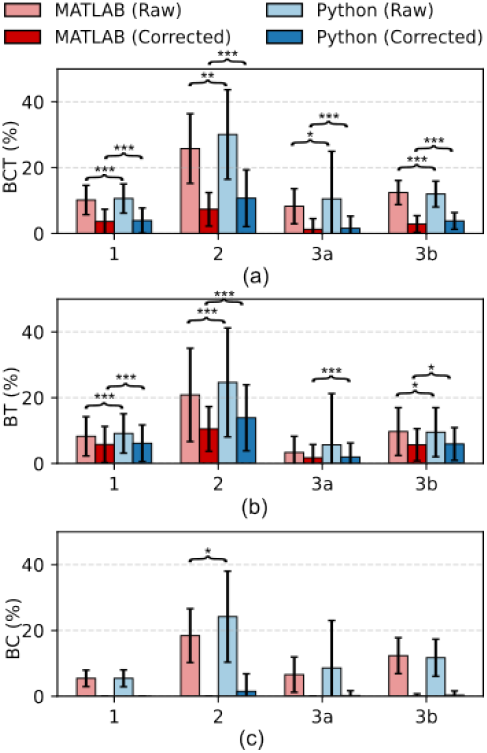
Comparison of artifact-detection and correction performance between the MATLAB and Python implementations of APICE. Panels show (a) BCT—percentage of bad data samples, (b) BT—percentage of bad time points, and (c) BC—percentage of bad channels across datasets. Dataset 1: sleeping neonates, 2: awake 5-mo-olds, 3a: freely moving 7-y-old children, 3b: freely moving adults.

It is important to note that no objective ground truth exists for determining which implementation performs “better” based on the detected and corrected artifacts. It remains unclear whether the MATLAB version—detecting fewer artifacts and therefore retaining more data—is more accurate, or whether the Python version—showing higher sensitivity to noise—is more effective at identifying subtle contamination. Without a definitive reference signal, these differences cannot be interpreted as superiority of one implementation over the other. Although APICE-Py was developed to closely mirror the original APICE implementation, software ecosystems inherently differ: as noted by Delorme (2023), MATLAB and Python toolboxes (e.g., EEGLAB versus MNE) implement core operations such as filtering differently, even when based on the same underlying principles. As a result, small numerical and algorithmic discrepancies between implementations are expected.

### D. Standardized Measurement Error (SME)

While data retention provides a general indication on how much signal is preserved after preprocessing, it does not necessarily capture the quality or reliability of the retained data. For instance, high retention rate could simply reflect minimal subject movement during recording rather than improved data quality, whereas low retention rate could correspond to cleaner and more reliable EEG signals. Therefore, to evaluate the quality of the retained EEG data, we compared the SME values between APICE and APICE-Py across Dataset 1 and Dataset 2 (Figure 4).

**Figure 4.**
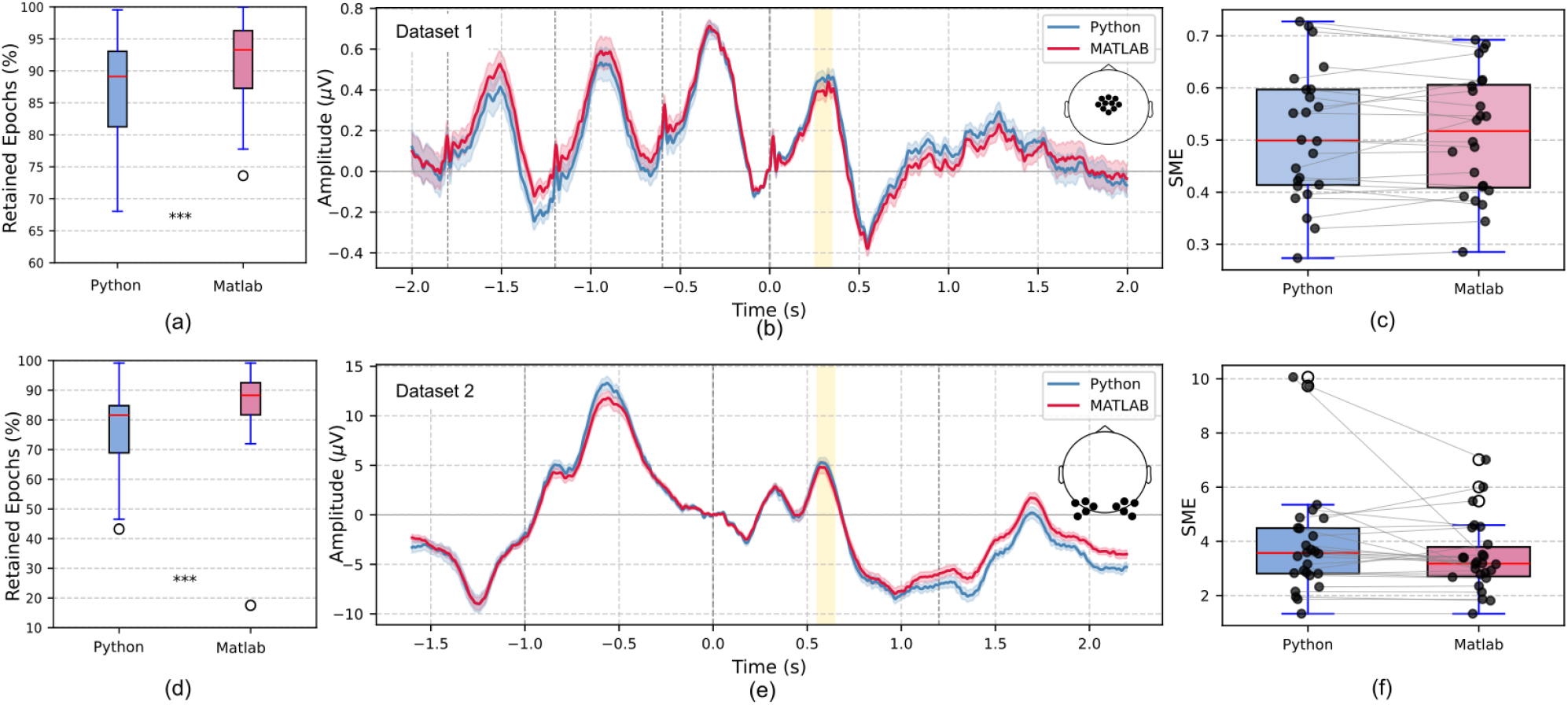
Comparison of the data quality between APICE and APICE-Py for Dataset 1 (auditory task with sleeping neonates) and Dataset 2 (visual task with 5-month old infants). Panels (a, d) show the number of retained epochs after preprocessing, (b,e) display the corresponding grand-average ERPs, and (c,f) present the SME values for each dataset.

We segmented the trials for Dataset 1 from -2 s to 2 s and for Dataset 2 from -1.6 to 2.2 s, applied both pipelines to the segments and rejected the bad epochs using the same criteria. As expected, APICE retained a higher percentage of epochs than APICE-Py (Fig. 4a,d), consistent with the Python version detecting more artifacts at the continuous-data level. Retention was significantly lower in APICE-Py for both datasets (Dataset 1: APICE-Py = 86.50 ± 8.22% vs. APICE = 90.84 ± 7.13%, p < 0.001; Dataset 2: APICE-Py = 75.57 ± 15.03% vs. APICE = 84.38 ± 15.16%, p < 0.001). We then computed the grand-average ERPs across all subjects at the regions of interest (ROIs): central electrodes for Dataset 1 (auditory task) and occipital electrodes for Dataset 2 (visual task). The ROIs are indicated on the electrode montages in Fig. 4. SME values were calculated within 250–350 ms for Dataset 1 to capture the auditory response, and within 550–560 ms for Dataset 2 to capture the P400 visual ERP component (Fló et al. (2022)). The ERP waveforms in Figure 4b,e show comparable average responses across the two implementations (Dataset 1: APICE-Py = 0.44 µV, APICE = 0.39 µV; Dataset 2: APICE-Py = 1.84 µV, APICE = 1.96 µV). Likewise, the SME values indicate no significant differences between the MATLAB and Python versions (Dataset 1: APICE-Py = 0.51 ± 0.12%, APICE = 0.51 ± 0.12%, p = 0.75; Dataset 2: APICE-Py = 3.92 ± 2.01%, APICE = 3.42 ± 1.28%, p = 0.09). These results suggest that APICE-Py performs comparably to the original MATLAB implementation in terms of data quality, despite detecting a slightly higher proportion of artifacts. In general, lower epoch retention can pose a concern for EEG studies, as fewer usable trials reduce statistical power and may increase variability in ERPs. However, the SME analysis showed no significant increase in measurement error for APICE-Py. It suggests that the additional segments flagged by APICE-Py were likely genuinely contaminated or of lower quality, and their exclusion did not compromise the reliability of the resulting ERPs.

### E. Signal-to-Noise Ratio (SNR)

Dataset 3 was recorded during a child–parent collaborative puzzle-solving task using a hyperscanning setup. Because this paradigm involves free movement and unconstrained social interaction, it introduces substantial motion artifacts and signal variability. Unlike stimulus-locked designs, hyperscanning studies rely on continuous measures such as inter-brain coherence and neural synchrony, which are highly sensitive to noise and fluctuations in signal amplitude. Poor SNR can artificially inflate or distort synchrony and connectivity estimates, leading to spurious interpretations of interpersonal neural coupling (Welke and Vessel (2022)). For these reasons, evaluating SNR provides an appropriate and meaningful assessment of data quality for hyper-scanning EEG.

We preprocessed the continuous signals (band-pass filtered between 0.2–40 Hz) and then computed the SNR across electrodes and subjects. Figure 5 compares the mean SNR values obtained from the two implementations. Overall, SNR values differed slightly across paired electrodes, with the Python implementation showing higher global SNR (Children: APICE-Py = 0.68 dB, APICE = –0.47 dB; Adults: APICE-Py = –0.29 dB, APICE = –1.84 dB), consistent with APICE-Py detecting and correcting a greater number of noisy segments. A particularly noticeable difference appears at the prefrontal electrodes (Fp1 and Fp2), which show substantially lower SNR in the MATLAB implementation. This discrepancy is expected because prefrontal sites are inherently more susceptible to ocular and facial-muscle artifacts (e.g., blinks, squinting, eyebrow movements), which occur frequently in naturalistic tasks. If APICE-Py identifies and corrects more of these frontal artifacts, it would naturally produce higher SNR in these channels. In addition, methodological differences between MNE-Python and EEGLAB/MATLAB—especially in filtering and spatial interpolation procedures (Delorme (2023))—may further account for the observed SNR differences at these highly artifact-prone sites. Overall, the SNR patterns indicate that discrepancies between the two implementations arise from the interaction of signal characteristics, artifact prevalence, and software-specific processing steps, rather than from systematic advantages of one pipeline over the other.

**Figure 5.**
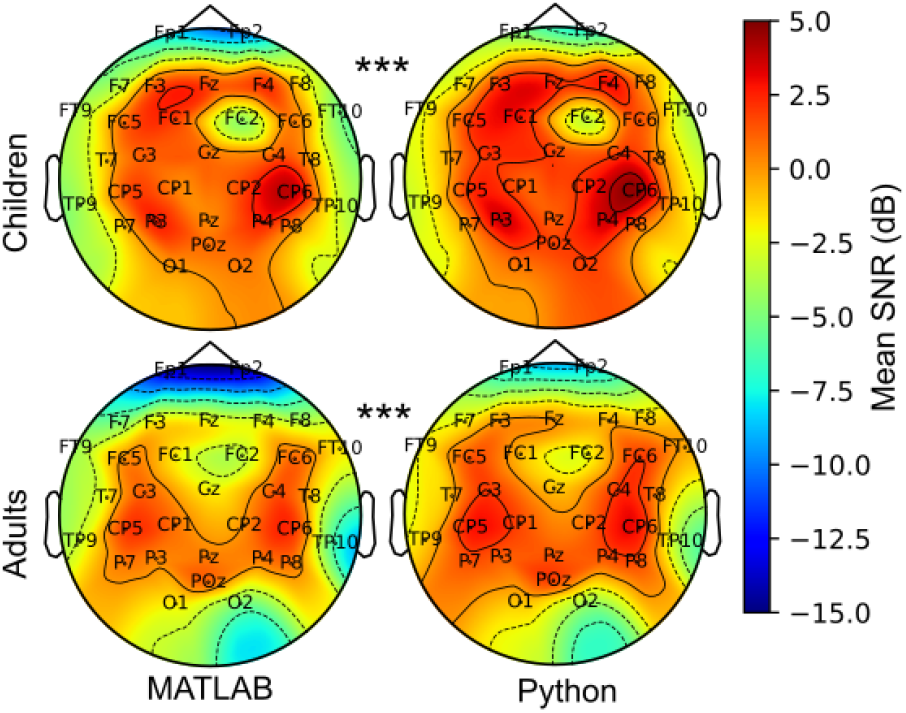
Signal-to-noise ratio (SNR) comparison between the MATLAB (APICE) and Python (APICE-Py) implementations for Dataset 3a (children) and Dataset 3b (adults).

### F. Computational Time

One concrete example of implementation-level differences concerns the interpolation procedures. Although both APICE and APICE-Py use Spherical Spline Interpolation (SSI), and both enforce the same criterion that a channel or segment is interpolated only when at least 50% of its neighboring electrodes are marked as good, the underlying software frameworks handle interpolation differently. In APICE-Py, we initially relied on MNE’s *interpolate*_*bads*() function to correct both segment-level and channel-level artifacts. However, during testing we observed that processing time increased substantially as the number of artifacts increased, as reflected in the MNE-Python boxplots in Figure 6. Detailed profiling revealed that MNE’s interpolation is computationally heavy because it operates directly on raw or epochs objects, which makes repeated interpolation steps particularly slow when the pipeline must loop over many short artifacted segments. This behavior contrasts with the MATLAB implementation, where interpolation is applied on simpler data structures and is therefore more lightweight. These differences illustrate how function-level design choices in Python and MATLAB can lead to noticeable discrepancies in processing time, even when the theoretical interpolation method (SSI) is the same.

**Figure 6.**
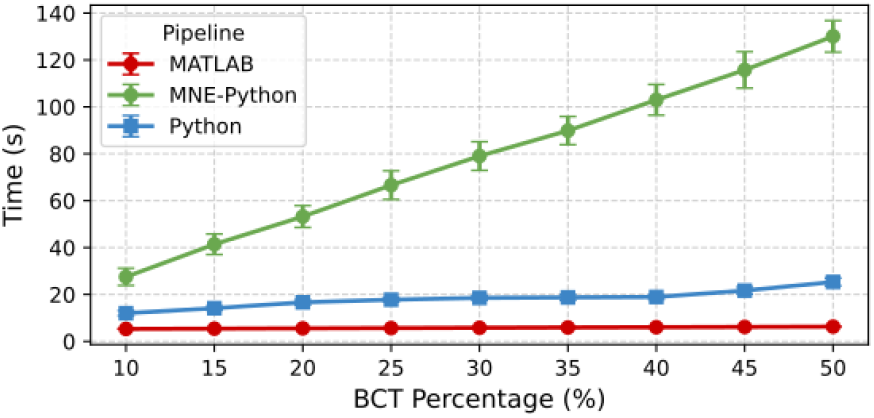
Comparison of computation time for MATLAB, MNE-Python, and customized APICE-Py SSI implementations across increasing levels of artifact load (% BCT).

We implemented a customized SSI function based on the original APICE (MATLAB) algorithm, and the computational bottleneck was further reduced by developing a parallelized version that bypasses the standard MNE data structures. This design avoids repeated object-level overhead, redundant data copying, and the cost of re-instantiating MNE objects at each iteration. In APICE-Py, this custom parallelized SSI function is used specifically for short-segment interpolation, where the number of interpolation calls is highest. For whole-channel interpolation—performed at least once per recording—APICE-Py still relies on MNE’s *interpolate*_*bads*(), as this step is less computationally taxing and maintains compatibility with MNE’s channel-handling conventions. This hybrid strategy substantially improves computational efficiency, as shown in Figure 6. When interpolation load increases (i.e., higher BCT percentages), the pure MNE-Python approach exhibits a steep, nearly linear rise in runtime, whereas the MATLAB version shows the lowest and most stable processing time. The customized APICE-Py implementation displays a much flatter slope, remaining computationally manageable even for heavily artifacted data.

Taken together, these results demonstrate that APICE-Py closely replicates the behavior of the original APICE pipeline while accommodating the practical differences inherent to the Python/MNE ecosystem. We also showed that despite APICE-Py detecting more artifacts, it still produced comparable ERP waveforms, SME values, and consistent SNR patterns across different datasets. Overall, the results support APICE-Py as a reliable and flexible preprocessing solution suitable for developmental to adult groups, controlled to naturalistic experimental paradigms, and various electrode configurations.

## Conclusion

In this work, we presented APICE-Py, an open-source, scalable, and reproducible preprocessing environment that extends the original APICE/MATLAB framework into the MNE-Python ecosystem. Its adaptive artifact-detection algorithms, hierarchical correction methods, and comprehensive quality reporting ensure consistent processing across datasets while maintaining transparency in all preprocessing decisions. Although APICE was originally developed for developmental EEG recorded with high-density wet electrode systems, we showed that both APICE and APICE-Py perform robustly across a broad range of EEG recordings—from sleeping neonates to 5-month-old infants, school-age children, and adults—spanning high-density laboratory EEG to low-density gel-based mobile systems, and from controlled experiments to naturalistic hyperscanning paradigms. Benchmarking against the validated MATLAB implementation demonstrated comparable data quality and reliability, as reflected in ERP responses, SME values, and SNR characteristics. The modular design of APICE-Py facilitates future extensions, including the addition of new artifact-detection strategies, support for emerging EEG hardware platforms, or integration with BIDS-compliant preprocessing workflows. Moreover, because the core of the pipeline relies on adaptive, data-driven thresholding rather than system-specific heuristics, similar strategies may be adapted for related modalities such as MEG, where continuous artifact detection and channel-level variability also pose significant challenges. Overall, APICE-Py provides a flexible and transparent framework that can accommodate the growing diversity of EEG research settings while promoting reproducibility and open science. Lastly, we emphasize that APICE-Py is not limited to cognitive neuroscience research; its flexibility makes it suitable for a wide range of EEG applications, including clinical assessments, educational and developmental studies, naturalistic and hyperscanning paradigms, and other EEG research domains.

## Acknowledgments

This work received funding from the European Research Council (ERC) under the European Union’s Horizon 2020 research and innovation program (grant agreement No. 695710 and 101142651) and from the French government managed by the Agence Nationale de la Recherche as part of the France 2030 program, under reference ANR-23-IAHU-0010.

This work was also funded by the French State through the Agence Nationale de la Recherche (grant ANR-21-ESRE-0034) under the Innovation, Data and Experiments in Education (IDEE) programme.

## Bibliography

Bigdely-Shamlo, N., Mullen, T., Kothe, C., Su, K.-M., and Robbins, K. A. (2015). The PREP pipeline: standardized preprocessing for large-scale EEG analysis. Frontiers in Neuroinformatics, 9:16. doi: 10.3389/fninf.2015.00016.

Buzzell, G. A., Morales, S., Valadez, E. A., Hunnius, S., and Fox, N. A. (2023). Maximizing the potential of eeg as a developmental neuroscience tool. Developmental Cognitive Neuroscience, 60:101201.

Chaddad, A., Wu, Y., Kateb, R., and Bouridane, A. (2023). Electroencephalography signal processing: A comprehensive review and analysis of methods and techniques. Sensors, 23(14):6434.

Coelli, S., Calcagno, A., Cassani, C. M., Temporiti, F., Reali, P., Gatti, R., et al. (2024). Selecting methods for a modular eeg pre-processing pipeline: An objective comparison. Biomedical Signal Processing and Control, 90:105830.

Delorme, A. (2023). Eeg is better left alone. Scientific reports, 13(1):2372.

Fló, A., Gennari, G., Benjamin, L., and Dehaene-Lambertz, G. (2022). Automated pipeline for infants continuous eeg (apice): A flexible pipeline for developmental cognitive studies. Developmental Cognitive Neuroscience, 54:101077.

Gramfort, A., Luessi, M., Larson, E., Engemann, D. A., Strohmeier, D., Brodbeck, C., et al. (2013). Meg and eeg data analysis with mne-python. Frontiers in Neuroinformatics, 7: 267.

Hervé, E., Mento, G., Desnous, B., and François, C. (2022). Challenges and new perspectives of developmental cognitive eeg studies. NeuroImage, 260:119508.

Kumaravel, V. P., Farella, E., Parise, E., and Buiatti, M. (2022). Near: An artifact removal pipeline for human newborn eeg data. Developmental Cognitive Neuroscience, 54:101068.

Luck, S. J. An introduction to the event-related potential technique. MIT press, (2014).

Luck, S. J., Stewart, A. X., Simmons, A. M., and Rhemtulla, M. (2021). Standardized measurement error: A universal metric of data quality for averaged event-related potentials. Psychophysiology, 58(6):e13793.

Mumtaz, W., Rasheed, S., and Irfan, A. (2021). Review of challenges associated with the eeg artifact removal methods. Biomedical Signal Processing and Control, 68:102741.

Radüntz, T. (2018). Signal quality evaluation of emerging eeg devices. Frontiers in physiology, 9:98.

Shirk, S. D., McLaren, D. G., Bloomfield, J. S., Powers, A., Duffy, A., Mitchell, M. B., et al. (2017). Inter-rater reliability of preprocessing eeg data: Impact of subjective artifact removal on associative memory task erp results. Frontiers in neuroscience, 11:322.

Welke, D. and Vessel, E. A. (2022). Naturalistic viewing conditions can increase task engagement and aesthetic preference but have only minimal impact on eeg quality. NeuroImage, 256:119218.

